# An efficient simplified method for the generation of corneal epithelial cells from human pluripotent stem cells

**DOI:** 10.1101/2022.01.16.476543

**Authors:** Rodi Abdalkader, Ken-ichiro Kamei

**Author notes:** Corresponding authors (R.A), (K.K).

## Abstract

Corneal epithelial cells derived from human pluripotent stem cells (hPSCs) are an important cell source for preclinical models to test ophthalmic drugs. However, current differentiation protocols lack instructions regarding optimal culturing conditions, which hinders the quality of cells and limits scale-up. Here, we introduce a simplified small molecule-based corneal induction method (SSM-CI) to generate corneal epithelial cells from hPSCs. SSM-CI provides the advantage of minimizing cell culturing time using two defined culturing media containing TGF-β, and Wnt/β-catenin pathway inhibitors, and bFGF growth factor over 25 days. Compared to the conventional human corneal epithelial cell line (HCE-T) and human primary corneal epithelial cells (hPCEpCs), corneal epithelial cells generated by SSM-CI are well-differentiated and express relevant maturation markers, including PAX6 and CK12. RNA-seq analysis indicated the faithful differentiation of hPSCs into corneal epithelia, with significant upregulation of corneal progenitor and adult corneal epithelial phenotypes. Furthermore, despite the initial inhibition of TGF-β and Wnt/β-catenin, upregulation of these pathway-related transcripts was observed in the later stages, indicating their necessity in the generation of mature corneal epithelial cells. Moreover, we observed a shift in gene signatures associated with the metabolic characteristics of mature corneal epithelial cells, involving a decrease in glycolysis- and an increase in fatty acid oxidation. This was also attributed to the overexpression of metabolic enzymes and transporter-related transcripts responsible for fatty acid metabolism. Thus, SSM-CI provides a comprehensive method for the generation of functional corneal epithelial cells for use in preclinical models.

## 1. Introduction

The corneal epithelium is a clear, light-permeable barrier that is critical in regulating homeostasis in the human eye, and is considered a major barrier for ocular drug delivery, representing the direct gate into the eye. Therefore, the corneal epithelium is an important determinant of drug absorption, distribution, metabolism, and excretion (ADME), as well as *in situ* toxicity, which must be bypassed during preclinical drug development [1].

A serious hurdle in the development of *in vitro* corneal epithelium models is the selection of the cell source. Inappropriate selection of cells can result in an unfaithful *in vitro* model, hampering the relevance of experimentation to human physiological conditions. Although several corneal cell lines, including SIRC1[2] and HCE-T [3][1], have played a critical role in the advancement of preclinical models of the cornea, interpreting data generated from cell lines is causing concerns among researchers. This is due to factors, such as cell contamination and the lack of expression of genes with relevant functions in cell characteristics [4][5]. Therefore, primary cells have gained renewed interest. However, obtaining primary corneal epithelial cells is not common practice [6] and is ethically unfavorable [7], while scientific analysis has shown that such cells have inter-individual differences[8], decline in biological functions after several passages [9], and are expensive to collect. Considering these issues, there is an urgent need for an alternative source of functional human corneal cells that is characterized by its biological relevance to the corneal epithelium, as well as the ability to grow and expand in *in vitro* cell culture systems.

Human pluripotent stem cells (hPSCs) are an important source of corneal cell lineages derived from embryonic stem cells (ESCs) or induced pluripotent stem cells (iPSCs). Protocols to induce the growth of eye tissue from hPSCs, such as the self-formed ectodermal autonomous multi-zone (SEAM) [10] and small molecule-based differentiation (SMD) methods have previously been reported [11][12]. The first method relies on the autonomous formation of eye tissues, which mimics the eye development process. The latter relies on the employment of small chemicals that inhibit TGF-β and canonical Wnt pathways in combination with human basic fibroblast growth factor (bFGF) to divert the cells towards the corneal epithelium lineage [13]. Compared to the SEAM, the SMD method has several advantages, including the use of xenobiotic- and serum-free culturing conditions; however, this method also has several serious limitations, such as the use of cell aggregates, which can affect the robustness of cell generation and the employment of uncharacterized culturing media, and can undermine the scale-up application. Moreover, the mechanism and key biological pathways in which the SMD method directs cells towards the corneal lineage and enhances the generation of adult corneal epithelial cells is not clearly known. Thus, the development of a simple yet efficient method for the generation of corneal epithelial cells in a short period using well-defined serum-free xeno-free cell culture conditions will provide a robust methodology to boost the generation of hPSC-derived corneal epithelial cells.

Herein, we have developed a simplified, small molecule-based corneal induction method (SSM-CI), with the following advantages: 1. An optimized hPSC culture method using a suspension expansion culture protocol. 2. Chemically defined culture medium free of xenobiotics and serum components. 3. Optimized concentrations of chemical inhibitors and growth factor bFGF. 4. Determination of key biological pathways controlling corneal lineage diversion. The potential of SSM-CI method was evaluated by the investigation of both eye developmental markers and corneal epithelial lineage markers, as compared to the conventional corneal epithelial cell line. Moreover, we conducted RNA sequencing (RNA-seq) analysis to highlight the critical involvement of several biological pathways that shape the cell lineage characteristics, including alterations in the metabolic features of cells, as well as the enzyme activities and small molecule transport.

## 2. Materials and Methods

### 2.1. Differentiation of corneal epithelial cells from human pluripotent stem cells

The human embryonic stem cells KhES1(K1), KhES1-OCT4-EGFP (K1-OCT4-EGFP) [14], and human induced pluripotent stem cells 585A1 were used following the guidelines of the ethics committee of Kyoto University. Prior to inducing differentiation, the cell culture dish was coated with Matrigel overnight at 4 °C. Cultured cells were washed with PBS and treated with TryPLE Express (Thermo Fisher Scientific, Inc., Waltham, MA, USA) at 37 °C for 5 min, followed by the addition of E8 medium (Thermo Fisher Scientific, Inc., Waltham, MA, USA) and transfer of the cell suspension into a 15-mL tube. Cells were centrifuged at 200 × *g* for 3 min, and the supernatant was removed and resuspended in E8 basal medium supplemented with 10 μM Y27632 (Wako, Osaka, Japan), and plated at a density of 1×10^4^ cell/cm^2^ on a Matrigel (Corning, Corning, NY, USA)-coated culture dish, and cultured for 24 h in a humidified incubator at 37 °C with 5% CO_2_. The culture medium was replaced with E8 basal medium daily. When cells reached 30-40% confluency, the E8 medium was replaced with chemical induction medium, comprising fresh E6 medium (Thermo Fisher Scientific, Inc., Waltham, MA, USA) supplemented with a Wnt pathway inhibitor (IWR-1 endo) (Selleck), TGF-β kinase/activin receptor-like kinase inhibitor (A83-01) (Wako, Osaka, Japan), and human bFGF (Wako, Osaka, Japan). These conditions were applied for four days. Subsequently, cells were cultured in maintenance medium comprising E6 medium alone (Table. S1). The medium was changed daily.

### 2.2. Human corneal epithelial cell culture

SV40-Adeno vector-transformed human corneal epithelial cells (HCE-T cells; RRID: CVCL_1272) were provided by the RIKEN Bioresource Research Centre (Ibraki, Japan). Cells were cultured in DMEM/F12 (Thermo Fisher Scientific, Inc., Waltham, MA, USA) supplemented with 5% (v/v) fetal bovine serum (Cell Culture Bioscience, Tokyo, Japan), 5 μg mL^-1^ insulin (Wako, Osaka, Japan), 10 ng mL^-1^ human epithelial growth factor (Nacalai), and 0.5% dimethyl sulfoxide (Wako, Osaka, Japan). The cells were passaged with trypsin-EDTA (0.25–0.02%) solution at a 1:4 subculture ratio. Primary Corneal Epithelial Cells: Normal, Human (ATCC^®^ PCS-700-010^™^) HCECs were obtained from the American Type Culture Collection (Lot#80915170, ATCC; VA, USA) and cultured in corneal epithelial cell basal medium containing the growth supplements Apo-transferrin (5 μg mL^-1^, epinephrine 1 μM, extract P (0.4%), hydrocortisone, Hemisuccinate 100 ng mL^-1^), L-glutamine (6 mM), insulin (5 μg mL^- 1^), and growth factor proprietary formulation (CE). The cells were passaged with trypsin-EDTA (0.05%–0.02%) at a density of ≥ 5000 cells per cm^2^. Cells in passage ≤ 5 were used in all experiments.

### 2.3. Immunofluorescence and microscopy imaging

For immunostaining, cells were fixed with 4% paraformaldehyde (Wako, Osaka, Japan) in PBS for 25 min at 25 °C, and then permeabilized with 0.5% Triton X-100 (MP Biomedicals, CA, USA) in PBS for 10 min at 25 °C. Subsequently, cells were blocked with blocking buffer containing 5% (v/v) normal goat serum (Vector), 5% (v/v) normal donkey serum (Wako, Osaka, Japan), 3% (w/v) bovine serum albumin (Sigma-Aldrich), and 0.1% (v/v) Tween-20 (Nacalai) at 4°C for 24 h, and then incubated at 4°C overnight with the primary antibody (Table. S2) in blocking buffer. The cells were then washed and incubated at 37°C for 60 min with the appropriate secondary antibody (Alexa Fluor 488 donkey anti-rabbit IgG and Alexa Fluor 594 donkey anti-mouse IgG 1:1000; Jackson ImmunoResearch, West Grove, PA, USA) in blocking buffer prior to a final incubation with 4,6-diamidino-2-phenylindole (DAPI) (Wako, Osaka, Japan) at 25°C. For imaging, we used a Nikon ECLIPSE Ti inverted fluorescence microscope equipped with a CFI plan fluor 10×/0.30 N.A. objective lens (Nikon, Tokyo, Japan), CCD camera (ORCA-R2; Hamamatsu Photonics, Hamamatsu City, Japan), mercury lamp (Intensilight; Nikon), XYZ automated stage (Ti-S-ER motorized stage with encoders; Nikon), and filter cubes for fluorescence channels (DAPI, GFP, and CY5; Nikon). Images were then analyzed using the ImageJ software (National Institutes of Health, Maryland, USA). Cell Profiler software (Version 3.1.8; Broad Institute of Harvard and MIT, USA) was used for analysis [15].

### 2.4. RNA extraction and sequencing

RNA was extracted using the RNeasy Mini Kit (Qiagen, Hilden, Germany) according to the manufacturer’s instructions. Sample quality was determined using a bioanalyzer (Agilent Technologies, Inc., USA) with, all samples verified to have an RNA integrity number (RIN) value ≥ 7. Samples were then subjected to next-generation sequencing (Macrogen, Tokyo, Japan) beginning with an RNA-seq library construction (TruSeq stranded mRNA LT Sample Prep Kit), followed by sequencing using the NovaSeq 6000 Illumina system with 101 reads per specimen.

### 2.5. RNA-seq data mining and GO enrichment analysis

Quality control of the sequenced raw reads was performed by calculating the quality of reads, total bases, total reads, GC (%), and basic statistics (FastQC v0.11.7). To reduce bias in the analysis, artifacts such as low-quality reads, adaptor sequences, contaminant DNA, or PCR duplicates were removed (Trimmomatic0.38)[16]. Trimmed reads were mapped to the reference genome using the splice-aware aligner HISAT2 (HISAT2 version 2.1.0, Bowtie2 2.3.5.1)[17][18]. The transcript was assembled using StringTie with aligned reads (StringTie version 2.1.3b) [19]. This process provides information on known, novel, and alternative splicing transcripts. Expression profiles were represented as read counts and normalization values, based on transcript length and depth of coverage. The fragments per kilobase of transcript per million mapped reads (FPKM) value or the reads per kilobase of transcript per million mapped reads (RPKM) was used as the normalization value. In groups with different conditions, differentially expressed genes or transcripts were filtered out through statistical hypothesis testing. The enrichment test was conducted with a significant gene list using the Profiler tool platform [20].

### 2.6. Statistical analysis and data visualization

The unpaired t-test and Tukey’s HSD comparison test were performed using Python Jupyter notebook 6.1.4, with Pandas and Bioinfokit packages [21]. Data visualization was performed using Python 3 with matplotlib and seaborn packages.

## 3. Results

### 3.1. hPSC culturing using the suspension expansion culturing method

To improve the robustness of the cell differentiation process, we used the suspension expansion culture method to passage hPSCs [22]. Single cells were collected on TrypleE and passed in xeno-free/serum-free E8, which is well-known to preserve the self-renewal properties of hPSCs [23]. hPSC colonies exhibited a unified and homogenous morphology with a defined edge, indicating their high quality [24].

### 3.2. SSM-CI to induce differentiation of hPSCs into corneal epithelial cells

As the diversion of corneal cells requires the inhibition of the TGF-β and Wnt/β-catenin signaling pathways, we employed a xeno-/serum-free E6 basal medium, which has the same composition as E8 media, except for the lack of a TGF-β inhibitor and bFGF. This medium allows the separate inclusion or absence of these components, without compromising the presence of other components which are necessary for proliferation and expansion of corneal cells during the differentiation process [25].

To direct hPSCs towards the corneal lineage, we used the IWR-1 endo Wnt/β-catenin inhibitor, A83-01 TGF-β kinase/activin receptor-like kinase inhibitor, and human bFGF to activate the FGF pathway (Fig. 1A). Initially, to determine the optimal conditions of chemical inhibitors and their effect on the promotion of hPSCs toward the corneal lineage, we tested the impact of different concentrations of IWR-1 endo and A83-01. K1 hESCs with enhanced green fluorescent protein (EGFP) expression driven by the *OCT4* promoter (K1-OCT4-EGFP) were treated with IWR-1 endo and A83-01 at concentrations of 2.5 and 10 μM, respectively, with or without 50 ng mL^-1^ bFGF for 4 days. Chemical induction with 10 μM IWR-1 endo and A83-01 led to the detachment of cell aggregates on day 1 (Fig. S1A), whereas treatment with 2.5 μM of IWR-1 endo and A83-01 without bFGF led to a low expression of PAX6 at day 7, despite the significant reduction in EGFP in K1-OCT4-EGFP (Fig. S1B). Conversely, induction with 2.5 μM of IWR-1 endo, 2.5 μM of A83-01, and 50 ng mL^−1^ of bFGF resulted in a homogenous cell morphology, with a significantly reduced EGFP signal by day 4, and no further changes noticed at day 8 (Fig. 1A, B). Under these conditions, cells expressed PAX6 on day 7 significantly more highly than the human corneal epithelial cell line HCE-T (Fig. 1C, D). At this point, the cell morphology changed to an epithelial-like polygonal morphology.

**Fig. 1.**
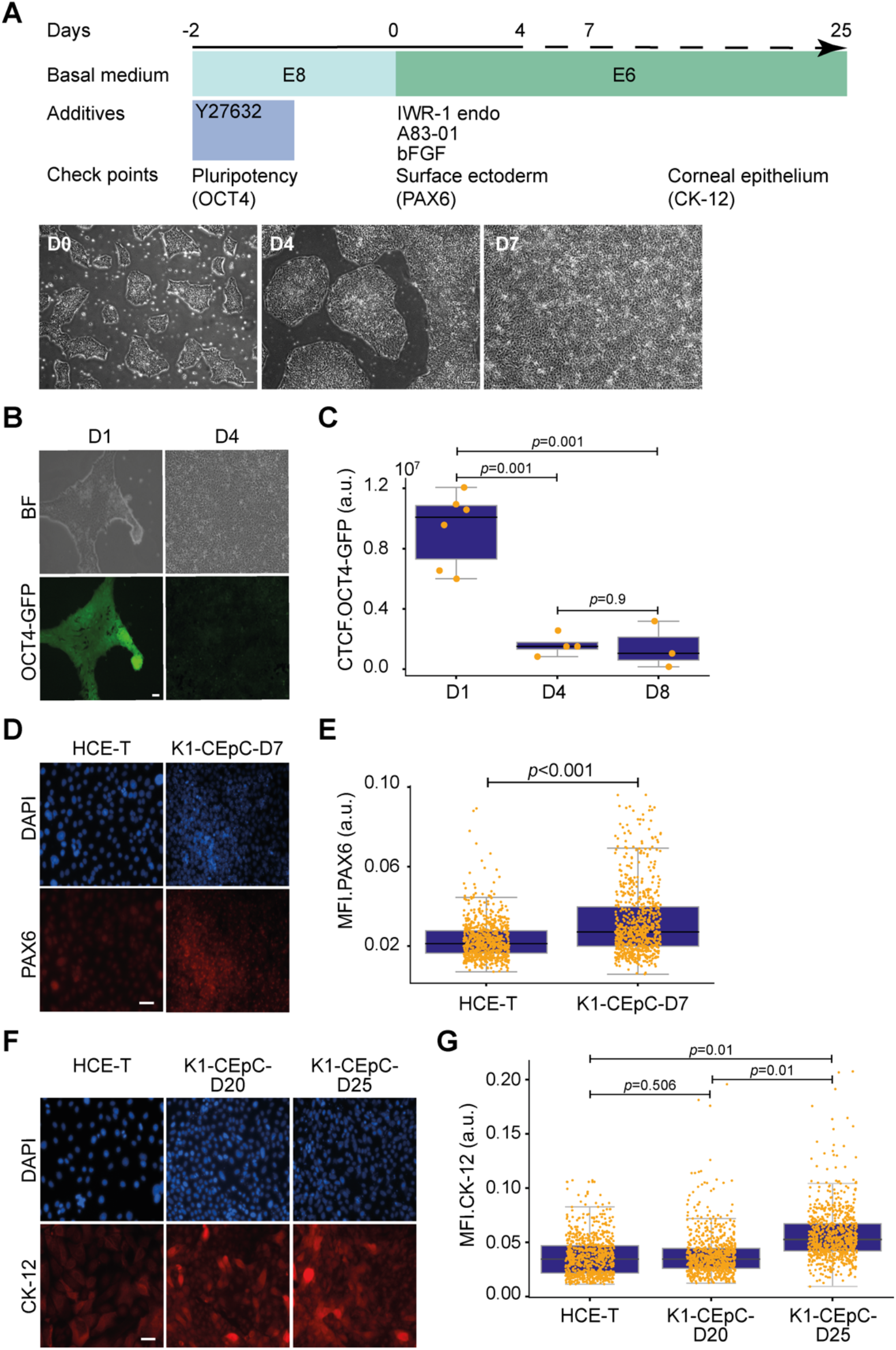
Generation of corneal epithelial cells form hPSCs. (A) An illustration of the overall protocol included a bright-field images of K1-CEpCs at D0, D4, and D7. Scale bar, 100 μm. (B) Fluorescent micrograph images of K1-OCT4-EGFP at D1, and D4. Scale bar, 50 μm. (C) Corrected total cell fluorescence (CTCF) of K1-OCT4-EGFP cells at D1, D4, and D8. (D) Fluorescent micrograph images indicating the expression of eye developmental marker (PAX6). Scale bar, 50 μm. (E) Single cells immunofluorescence analysis of PAX6. (F) Fluorescent micrograph images indicating the expression of corneal epithelium maturation marker (CK-12). Scale bar, 50 μm. (G) Single cells immunofluorescence analysis of CK12. Boxplot in which the median of each group is indicated with a black line (25^th^ to 75^th^ interquartile range). The *p*-values were determined using Tukey HSD’s multiple comparison test as well as the unpaired t.test.

Considering that the E6 basal medium contains all of the components required to induce corneal cell maturation, including essential amino acids, insulin, and transferrin [26], this medium was further utilized to induce cell maturation. We subsequently investigated the commitment and maturity of corneal epithelial cells by testing the expression of the corneal epithelial maturation marker cytokeratin 12 (CK12). We noticed a clear upregulation of CK12 on day 20, which was further boosted by day 25 compared to in HCE-T cells (Fig. 1E, F). Application of the same protocol to 585A1 iPS cells led to a CK12 positivity percentage of 56.1±18.2 % at day 20 versus 73.1±18.6% in K1-OCT4-EGFP cells (Fig. S2A, B).

### 3.3. RNA-seq analysis of transcripts and related pathway in hPSC-CEpCs

To investigate the temporal transcriptome signature of the biological pathways activated during the SSM-CI method-based differentiation, we selected K1 cells for RNA-seq analysis of the derived CEpCs on days 0 (D0), 10 (D10), and 20 (D20). Comparison of D10 vs. D0 revealed a total of 6879 differentially expressed genes (DEGs) identified on volcano plots, among which 3649 were upregulated genes and 3230 were downregulated (Fig. 2A).

**Fig. 2.**
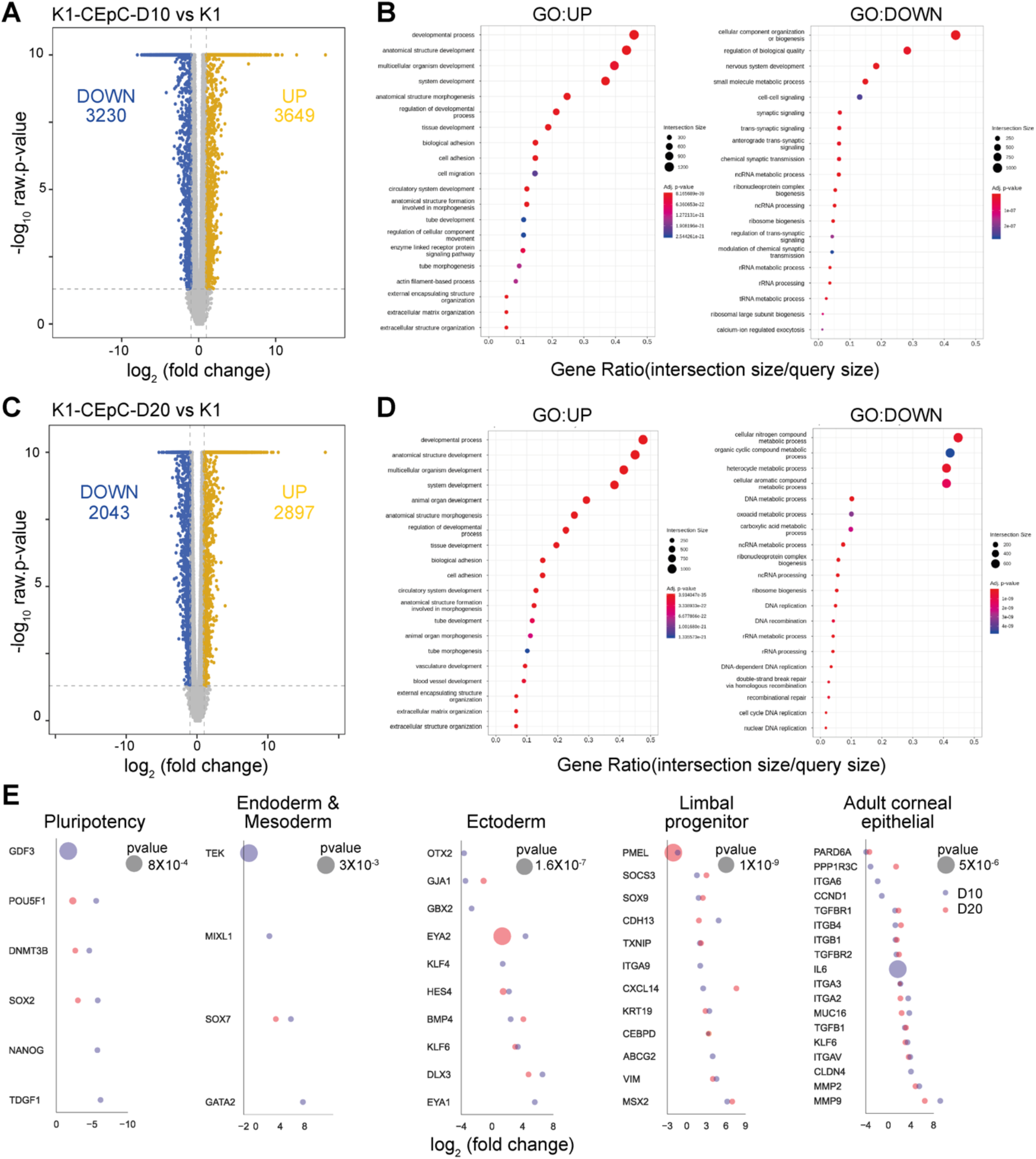
RNA-seq analysis showing the genes signatures and biological pathways during the diversion of cells into the corneal lineage. (A). The volcano plot representing the differentially expressed genes (DEGs) of K1-CEpCs at D10. Genes with 2-fold change and a p-value <0.05 was applied. Blue: downregulated genes, yellow: upregulated genes. (B). GO pathways of DEGs in K1-CEpCs at D10. (C) The volcano plot representing the DEGs of K1-CEpCs at D20. (D) GO pathways of DEGs in K1-CEpCs at D20. (E). DEGs of the pluripotency markers, endoderm and mesoderm markers, ectoderm markers, corneal progenitor markers, and adult corneal epithelial markers.

At D10, significantly upregulated genes were annotated with GO pathways such as tissue development (GO:0009888), cell adhesion (GO:0007155), enzyme-linked receptor protein signaling (GO:0007167), actin filament-based process (GO:0030029), extracellular matrix organization (GO:0030198), and response to transforming growth factor beta (GO:0071559). The downregulated genes annotated to several GO pathways, including nervous system development (GO:0007399) and small molecule metabolic process (GO:0044281 (Fig. 2B). At D20, 2897 upregulated and 2043 downregulated genes were annotated, corresponding to the previously mentioned pathways, as observed in D10 (Fig. 2C). Interestingly, downregulated genes were highly and specifically annotated to cellular nitrogen compound metabolism (GO:0034641), metabolic process of organic cyclic (GO:1901360)/ aromatic (GO:0006725)/ carboxylic (GO:0019752)/ oxoacid compounds (GO:0043436), as well as their involvement in DNA replication/cycle and processing (Fig. 2D).

To investigate the diversion of cells towards the corneal lineage, we examined the expression of specific genes (log2 fold differences ≥1). We thus assured the commitment towards the corneal lineage during the transition process from the pluripotency state toward one of the three germ layers (endoderm, mesoderm, ectoderm), as well as the emergence of corneal progenitor phenotypes and commitment towards differentiation into adult corneal cells (Fig. 2E).

At D10, there was a significant decrease in pluripotency markers, with *TDGF1, NANOG, SOX2, DNMT3B, OCT4* (*POU5F1*), and *GDF3*. At D20, *GDF3, OCT4* (*POU5F1*), *DNMT3B*, and *SOX2* still only lowly expressed. Among the endoderm and mesoderm markers, only an increase in *GATA2, SOX7*, and *MIXL1*, and a decrease in *TEK* were observed at D10. At D20, only *SOX7* was significantly upregulated. The ectoderm markers *EYA2, KLF4, HES4, BMP4, KLF6*, and *DLX3* were significantly upregulated at both D10 and D20. In contrast, there was a notable increase in the limbal progenitor gene markers including *MSZX2, VIM, ABCG2, CEBPD, KRT19, CXCL14, ITGA9, TXNIP, CDH13, SOX9*, and *SOCS3*, and a decrease in *PMEL* at D10. At D20, all the aforementioned genes were upregulated, except for *ABCG2*. Among the 18 gene markers of adult corneal epithelial cells, 14 were significantly upregulated at both D10 and D20, including *MMP9, MMP2, CLDN4, ITGAV, KLF6, TGFB1, MUC16, ITGA2, ITGA3, IL6, TGFBR2, ITGB1, ITGB4*, and *TGFBR1*. Conversely, *CCND1, ITGA6*, and *PARD6A* were downregulated at both timepoints, while *PPP1R3C* was downregulated at D10 but significantly upregulated at D20.

### 3.4. Cell characteristics and maturation ability in comparison with human primary corneal epithelial cells

To investigate the equivalent expression of generated cells, DEG analysis was conducted on embryonic stem cells (K1), human primary corneal epithelial cells (hPCEpC), and K1-CEpC at D20. The cluster analysis of gene expression represented in the dendrogram indicated the deviation of gene expression in both hPCEpC and K1-CEpC-D20 from K1, where hPCEpC and K1-CEpC-D20 had a lower height than K1 (Fig. 3A). To highlight the similarities and differences between hPCEpC and K1-CEpC at D20, we conducted a DEG analysis of hPCEpC versus K1-CEpC. We identified a total of 4717 DEGs, among which 1845 were upregulated and 2872 were downregulated (Fig. 3B). The top GO biological pathways of the upregulated genes were related to the response to stress (GO:0006950), regulation of multicellular organismal process (GO:0051239), programmed cell death (GO:0012501), biological adhesion (GO:0022610), and small molecule biosynthetic process (GO:0044283). The top GO biological pathways of the downregulated genes were associated with anatomical structure development (GO:0048856), cell adhesion (GO:0007155), cell morphogenesis (GO:0000902), and cell junction organization (GO:0034330) (Fig. 3C).

**Fig. 3.**
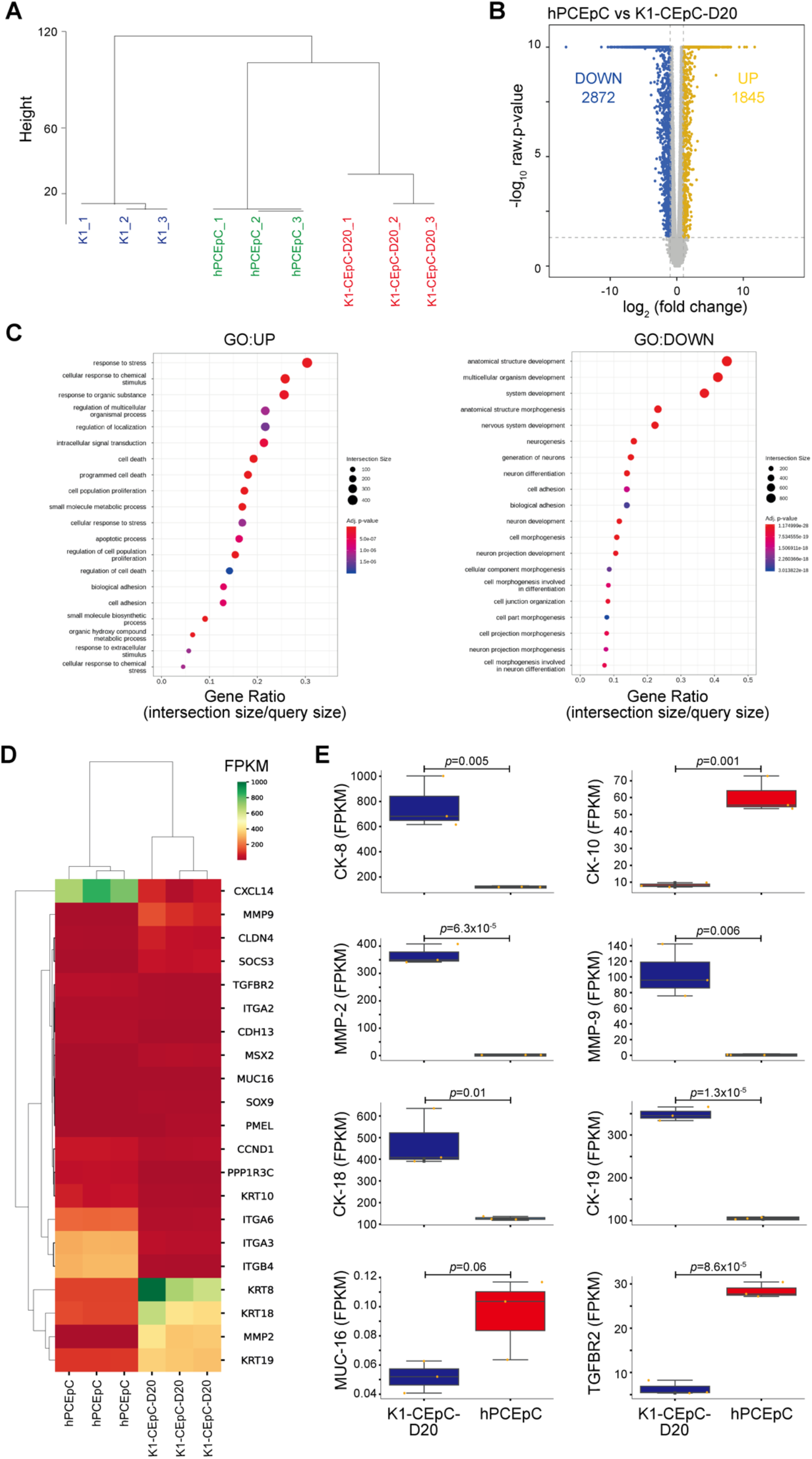
Maturation characteristics of K1-CEpCs versus hPCEpC represented by RNA-seq analysis. (A). Dendrogram indicating the hierarchical Clustering analysis by showing the expression similarities among K1, hPCEpC, and K1-CEpCs-D20. (Distance metric = Euclidean distance, Linkage method= Complete Linkage). (B) The volcano plot representing the differentially expressed genes (DEGs) of hPCEpC vs K1-CEpCs at D20. Genes with 2-fold change and a p-value <0.05 was applied. Blue: downregulated genes, yellow: upregulated genes. (C) GO pathways of DEGs in hPCEpC vs K1-CEpCs-D20. (D) Heatmap of selected genes related to adult corneal epithelium markers in hPCEpC and K1-CEpCs-D20. (E) The comparative gene expression (normalized FPKM values) of essential adult corneal epithelial markers in hPCEpC vs K1-CEpCs at D20. Boxplot in which the median of each group is indicated with a black line (25^th^ to 75^th^ interquartile range). The *p*-values were determined using the unpaired t.test.

We further compared the levels of gene expression (normalized FPKM values) of 21 essential adult corneal epithelial gene markers (Fig. 3D). Although expression of these genes was observed in both hPCEpC and K1-CEpC-D20, we noticed a significant upregulation of *CK-8*/*CK-18/CK-19* and downregulation of *CK-10* in K1-CEpC as compared to hPCEpC. Moreover, there was a significant upregulation in expression of the metalloprotease family members *MMP2* and *MMP9*, which are known as functional markers of the corneal epithelium, while there was a lower expression of the glycoprotein mucin16 (*MUC 16*) and *TGFBR*2 K1-CEpC than in hPCEpC (Fig. 3E).

### 3.5. Significant upregulation of TGF-ß and WNT/ ß-catenin pathways during corneal lineage diversion

To further investigate other signaling pathways contributing to the TGF-β and Wnt/ β-catenin pathways, we selected genes of the top GO biological pathways related to upregulated genes at D10 and D20. We observed significant involvement of genes in the TGF-β family member proteins, including SMAD activation, signaling by receptor tyrosine kinases, signaling by BMP, insulin growth factor signaling, signaling by NODAL, β-catenin in dependent on WNT signaling, transcriptional regulation by RUNX2, and transcriptional activities by *SMAD2*/ *SMAD3*/ *SMAD4*. In fact, among the SMAD transcripts, we observed a notable expression of *SMAD3*, a member of the TGF-β/Nodal/actin signaling transduction pathway, at both D10 and D20. SMAD transcripts acting in BMP/GDF signaling transduction were also notably expressed, including *SMAD6* at D10 and *SMAD7*/*9* at D10 and D20. *SMAD6, SMAD7*, and *SMAD9* were expressed in hPCEpC but at lower levels that K1-CEpCs-D20 (Fig. S3A). Moreover, we observed upregulation of members of the WNT receptor family, including *WNT3, WNT4, WNT5A, WNT5B, WNT7B, WNT9B*, and *WNT10B* at D10 and D20, while *WNT8B* was significantly downregulated at D10.

*WNTB, WNT10B*, and *WNT3* were equivalently expressed in hPCEpC and K1-CEpCs-D20, while *WNT5A, WNT9A*, and *WNT5B* were more strongly expressed in K1-CEpCs-D20 (Fig. S3B). Correspondingly, we observed an upregulation in TGF-β family proteins including *TGFB1/ TGFB3* (D10) and *TGFB2* (D10, D20). The TGF-β receptor family members, such as *TGFR1/TGFR2/TGFR3*, were significantly increased at both D10 and D20, while *TGFBR3L* expression was downregulated (Supplementary Fig. 4c). (Supplementary Fig. 4d). Compared to hPCEpC, there was no difference in the expression of *TGFB2OT1/ TGFB3* at D20, but rather a slight increase in *TGFB21/TGFB1I1/TGFR3* in K1-CEpCs, while *TGFR3* was more prevalent in hPCEpCs (Fig. S3C).

### 3.6. Gene signatures associated with metabolic features of cells during corneal lineage diversion

As we noticed a significant relationship between genes related to metabolic activity in the GO pathways, we investigated the metabolic gene signatures of the derived corneal cells during the differentiation process. Thus, we investigated changes in the expression of genes related to the glycolysis pathway, a known feature of hPSCs [27][28], and oxidative phosphorylation/fatty acid oxidation, both metabolic pathways known to be important in regulating the lineage of differentiated cells [29]. As expected, we identified only a few upregulated genes related to glycolysis. Conversely, we found more upregulated genes related to oxidative phosphorylation/fatty acid oxidation metabolism, indicated that the metabolic status of cells differentiating from a pluripotent state into the corneal lineage was drastically changed (Fig. 4A). We further noticed a correlated decrease in genes related to glycolysis metabolism, including *GPI* and *ENO2*, in hPCEpC as compared to K1-CEpCs-D20 and K1 (Fig. 4B, C).

**Fig. 4.**
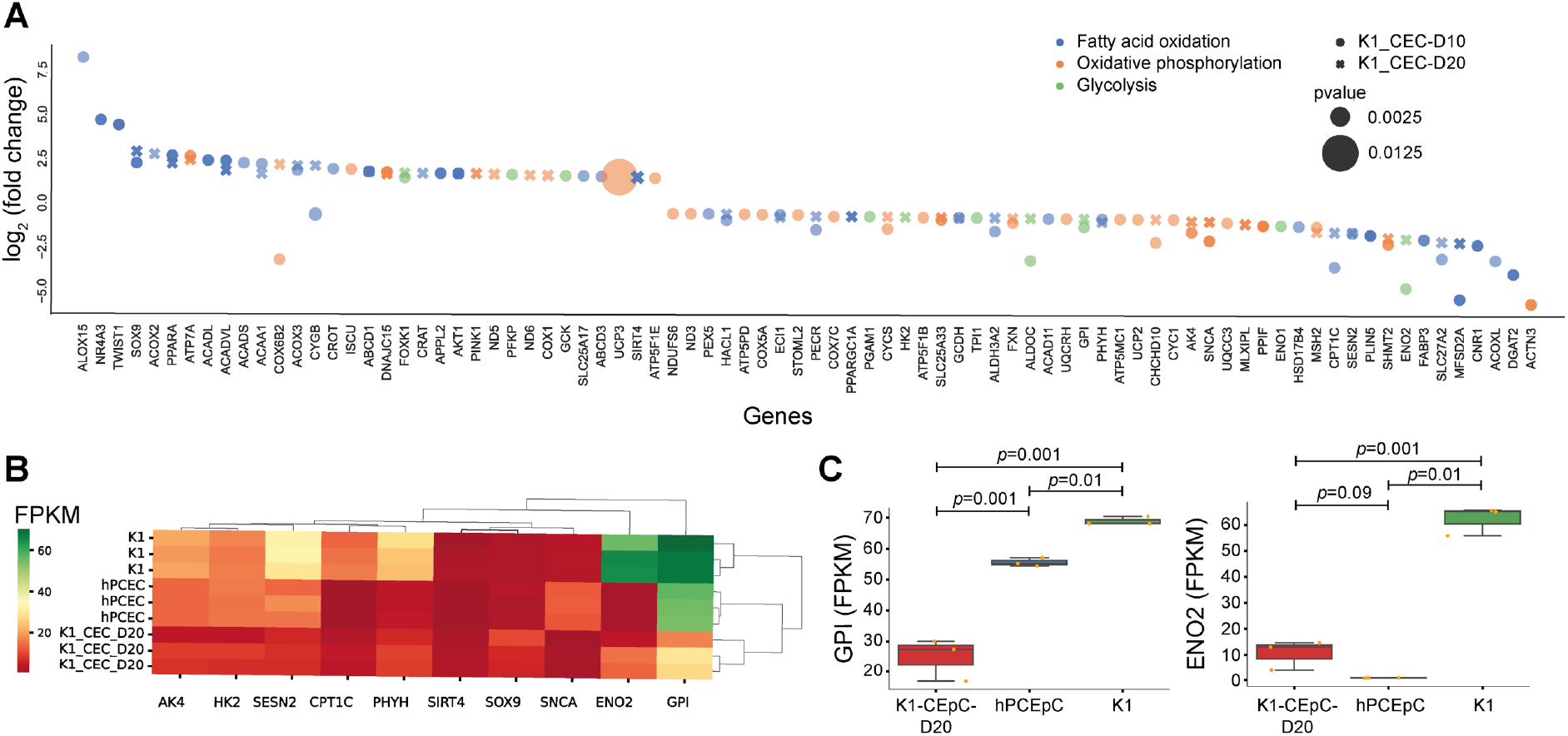
The alteration of the metabolic characteristics of K1-CEpCs represented by gene signatures associated with metabolic features of cells. (A). DEGs representing fatty acid oxidation (blue), oxidative phosphorylation (orange), and glycolysis (green) of in K1-CEpCs at D10 and D20. Genes with 2-fold change and a p-value <0.05 was applied. (B). Heatmap of selected genes related to fatty acid oxidation and glycolysis in K1, hPCEpC, and K1-CEpCs-D20. (C) Boxplots indicating the comparative gene expression (normalized FPKM values) of glycolysis markers in K1-CEpC-D20, hPCEpC, and K1. Boxplot in which the median of each group is indicated with a black line (25^th^ to 75^th^ interquartile range). The *p*-values were determined using the unpaired t.test.

In addition to the glycolysis pathway and oxidative phosphorylation/fatty acid oxidation, a variety of cytochrome P450 (CYP) family enzyme-related genes that are responsible for cholesterol and all trans retinoic acid metabolism were also significantly upregulated, including *CYP26B1* (D20), *CYP46A1* (D10), *CYP27C1* (D20), *CYP1B1* (D10), CYP2C8 (D10), and *CYP7B1* (D20, D20). A significant downregulation of genes related to metabolic enzymes for prostaglandin metabolism (*CYP2S1*), vitamin D (*CYP27B1*), and polysaturated fatty acids (*CYP2J2*) was observed in D10 and D20. There were no notable differences in the expression level of *CYP2J2/ CYP2T1P/ CYP27B1/ CYP27A1* in hPCEpC compared to K1-CEpCs-D20. However, there was a slightly higher expression of *CYP1B1* in hPCEpC (Fig. S4A).

### 3.7. Gene signatures associated with molecular transporters of cells during the corneal lineage diversion

Among ATP-binding cassette (ABC) transporter transcripts, significant upregulation of *ABCA1* and *ABCG2* was observed at D10. *ABCA1* is a transporter that is highly expressed in the retina and facilitates the transportation of N-retinyl-phosphatidylethanolamine and *ABCG2*, which are specific markers of corneal limbal stem cells, which plays an essential role in the transportation of retinoic acid. Interestingly, there was a clear upregulation of the two transporters responsible for trafficking cholesterol: *ABCA5* (D10, D20) and *ABCC2* (D20), for which we noticed an equivalent level of expression in hPCEpC (Fig. S4B).

Regarding solute carrier transporter (SLC)-related transcripts, we also observed a clear downregulation of glucose transporter 4 (*SLC2A4*), lysine transporter (*SLC15A3*), and glutamate transporters (*SLC1A2, SLC1A6*, and *SLC7A17*). The upregulation of two other subunits of glutamate transporters (*SLC1A1* and *SLC1A3*) and their level of expression was similar to that in hPCEpC. The zinc transporter (*SLC39A7*) and glucose transporter 3 (*SLC2A3*) were more highly expressed in K1-CEpCs-D20 as compared to hPCEpC. In contrast, glucose transporter 1 (*SLC2A1*) was predominantly expressed in hPCEpC (Fig. S4C).

## 4. Discussion

The derivation of corneal epithelial cells from hPSCs has significant advantages over cell lines in preclinical trials. Although there has been progress in the development of differentiation methods, such as the SMD method, using small molecule inhibitors in xenobiotic-free and serum-free culturing conditions [11][13], these methods still lack knowledge regarding the optimal culturing conditions, which can affect the robustness and the quality of cells as well as the scale-up application of cell production.

In this study, we developed a simplified small molecule-based corneal induction method (SSM-CI) for the generation of corneal epithelial cells from hPSCs, for which we have developed optimal cell culturing conditions, including: 1. the employment of a single-cell expansion culturing system of hPSCs, 2. The minimization of cell-culturing steps by dividing the differentiation protocol into three stages: pluripotency maintenance, chemical induction, and maturation stages, 3. The use of only two defined xenobiotic-free and serum-free culture media: E8 medium for the pluripotency maintenance stage and E6 medium for the chemical induction and maturation stages. Thus, unlike the previous methods, SSM-CI provides an optimized and efficient method for the generation of human corneal epithelial cells in a short duration of only 20-25 days.

Among the chemical inhibitors for Wnt/ β-catenin inhibition, IWP-2 and IWR-1 endo have been extensively reported in many differentiation protocols, involving neuron cells [30][31], cardiomyocytes [32], and retinal and corneal cells, where both compounds have an equivalent inhibition potency [33]. For the inhibition of TGF-β kinase/activin receptor, several compounds are commonly reported in the differentiation of the forehead and eye lineage, such as SB431542 [34], SB505124 [35], and A83-01 [33], where the inhibition potency of A83-01 is much greater than that of other TGF-β/R inhibitors [36]; thus, we selected A83-01 to be combined with IWR-1 endo and bFGF for the initiation of differentiation toward the corneal epithelium lineage. In this process, we noticed two important factors for the success of the protocol: 1. The concentration of IWR-1 endo and A83-01 was critical, where 10 μM led to cell aggregation and detachment at D1 while 2.5 μM treatment resulted in excellent morphology. 2. The presence of bFGF was essential, and the absence of bFGF led to reduced expression of PAX6. During eye development, ectodermal cells express PAX6 to generate ocular surface epithelia, and cells that do not express PAX6 consequently divert into the epidermis lineage [37][38]. Our results correspond to the literature, showing that bFGF signaling is required to maintain PAX6 expression in the eye tissue and is an essential factor for diverting cells towards the eye lineage [39].

Next, to investigate the development of cells into mature corneal epithelial cells, cells were maintained for 25 days in the basal medium E6. E6 is a defined basal medium containing all of the components incorporated in DMEM/ F12 medium with the addition of ascorbic acid, insulin, and transferrin, which are required for the growth and expansion of corneal epithelial cells [26]. Thus, we employed E6 as a bifunctional medium that could fit in both the chemical induction stage and the maturation stage. CK-12 promoter activity is linked to the overexpression of the transcription factor PAX6 [40][41]; therefore, CK-12 is expected to be expressed under optimal maturation conditions. We noticed strong expression of CK-12 in cells at D20 and D25, indicating the successful development of corneal progenitor cells into adult corneal epithelial cells in the E6 medium.

We subsequently conducted RNA-seq analysis of cells at both the early differentiation stage (D10) and the maturation stage (D20). GO analysis of upregulated genes showed that they were mainly related to tissue development, cell adhesion, actin filament-based process, and extracellular matrix organization, which implied the actual diversion of the cell phenotypes into different cellular lineages, and specifically the alteration towards the corneal progenitor lineage considering ECM remodeling, the upregulation in cell adhesion and actin components that contain essential markers for the corneal epithelium lineage such as *ITGB1, MMP9*, and cytokeratin family such as *CK-8, CK-18*, and *CK-19*. This was further supported by the observed significant upregulation of corneal lineage markers, the corneal progenitor markers such as *ABCG2*, and the downregulation of pluripotency markers, neuron cell markers, endoderm lineage markers, and mesoderm lineage markers.

We subsequently compared hPCEpC with K1-CEpC-D20 at D20. GO biological pathways of the upregulated genes were related to the response to stress, regulation of multicellular organismal processes, programmed cell death, biological adhesion, and small molecule biosynthetic processes. Moreover, the top GO biological pathways of the downregulated genes were associated with anatomical structure development, cell adhesion, cell morphogenesis, and cell junction organization, which are essential features in corneal epithelial cells, the imbalance of which diminishes the overall characteristics of the corneal epithelium; however, hPCEpC features can gradually be degraded by cell death during cell expansion. Hence, we compared the expression of essential adult corneal epithelial markers such as in the intermediate filament expression; *CK-8* and *CK-18* are known for their expression in the corneal limbal cells as well as in the mature corneal epithelial cells [42]; CK-19 is known for its expression in both the cornea and conjunctiva [43], while CK-10 is generally expressed in the epidermal epithelium, in the skin as well as in corneal keratinocytes [44]. We noticed a significant upregulation of *CK-8*/*CK-18*/*CK-19* in K1-CEpC-D20 compared to hPCEpC. Surprisingly, there was substantial expression of the keratinocyte marker *CK-10* in hPCEpC. Simultaneously, the level of *TGFBR2*, which is known to be highly expressed in the corneal stroma mainly from keratinocytes, was also significantly higher in hPCEpC, indicating the heterogeneous properties of hPCEpC cells, which potentially incorporate keratinocyte populations among the corneal epithelial cells. Moreover, it is known that the metalloprotease family members MMP2 and MMP9, as well as the mucin glycoprotein MUC16, are actively secreted in adult corneal epithelial cells [45][46]. The expression of *MMP2/MMP9* was superior in the K1-CEpC-D20, while there was a lower expression level of *MUC16* compared to hPCEpC. It has been reported that *MUC16* is highly upregulated following the formation of the corneal barrier during the stratification of cells [46]. Considering that K1-CEpC-D20 is grown in the liquid-liquid interface, lower secretion of MUC16 is expected compared to hPCEpC isolated from the fully stratified corneal barrier. Thus, our results confirmed the maturation characteristics and purity features of cells generated by SSM-CI.

It is known that the triple signaling activities of Wnt/β-catenin, TGF-β/Nodal, and BMP are necessary for the development and formation of the eye [47][11][13]. Even though IWR1 and A-83-01 were applied for four days, their inhibitory potency on cells was clearly reversed, as our data showed that a week post chemical induction, the expression of TGF-β receptors and their downstream cytoplasmic signaling molecules (Smad2 and Smad3) and the expression of Wnt/β-catenin family secreted proteins were upregulated at both D10 and D20. The Smad cascade pathway is uniquely specific to TGF-β/activin signaling, where receptor-activated Smad proteins, Smad2 or Smad3, are phosphorylated by the TGF-β type I receptor kinase (TGFβRI), in concert with the common mediator, Smad4, after which they translocate to the nucleus where they play an important role in the activation of TGF-β/Smad-dependent genes. Smad signaling has been shown to be critical in the recovery of wounds in many tissues, including the lens and retina of the eye [33]. To confirm the necessity of these pathways in the maturation stage of corneal epithelial cells, we compared the expression of TGF-β and Wnt/β-catenin signaling genes in K1-CEpC-D20 at D20 versus hPCEpC. We noticed upregulation of genes involved in the SMAD (*SMAD6/7/9*) and *WNT (WNT2B/WN5TB/ WN9TB/ WN10TB*) pathways in both K1-CEpCs and hPCEpC, that the later activation of the TGF-β and Wnt/β-catenin pathways is necessary for mature corneal epithelial cells.

GO analysis of downregulated genes showed that they are related to nervous system development, indicating that even if cells express ectoderm lineage makers, they are not necessarily committed to the neuron cell lineage. Moreover, the downregulated genes were annotated to metabolic processes, including organic cyclic/aromatic/carboxylic/oxoacid compounds and cellular nitrogen. Therefore, we sought to further investigate this by evaluating gene signatures associated with the metabolic signature of cells in the early (D10) and late (D20) stages of differentiation. Interestingly, we noticed a dominant upregulation of oxidative phosphorylation as well as fatty acid oxidation-related transcripts in differentiated cells and the downregulation of major genes related to glycolysis. This phenomenon is also known as the Warburg effect [48], where stem cells initially depend on the use of glycolysis metabolism to guarantee their self-renewal, losing dependency on this pathway following differentiation [49]. This has already been elucidated with other types of cell lineages, including hPSC-derived cardiomyocytes and neuron cells. In hPCEpC, there was a corresponding decrease in the expression of some essential glycolysis genes, such as GPI and ENO2. Cellular metabolism involves two main processes: metabolic enzymes and transportation, including the influx and efflux of nutrients. To this end, we investigated the CYP-related enzyme genes for phase I metabolism, as well as two main membrane transporters, the ABC transporter family, and the SLC-related transporters. CYP family enzymes responsible for cholesterol, steroid, and lipid metabolism, such as *CYP1A1* and *CYP27A1/A2* were significantly upregulated in both K1-CEpCs and hPCEpC. Moreover, among the ABC transporter transcripts, significant upregulation of *ABCA1* and *ABCG2* was observed at D10. *ABCA1* is a transporter that is highly expressed in the retina and facilitates the transportation of N-retinyl-phosphatidylethanolamine, and *ABCG2*, which is a specific marker for corneal limbal stem cells, while playing an essential role in the transport of retinoic acid. Interestingly, we observed a clear upregulation of the expression of two transporters responsible for cholesterol trafficking: *ABCA5* (D10, D20) and *ABCC2* (D20), for which we noticed an equivalent level of expression in hPCEpC. It is known that the SLC7 family of transporters contributes primarily to the transportation of fatty acids, specifically cholesterol [50], as we noticed upregulation in the expression levels of *SLC7A1/A2/A5/A7/A8* in both K1-CEpCs and hPCEpC. Furthermore, we observed overexpression of *SLC2A3, SLC7A5, SLC16A3*, and *SLC38A2*, as well as *ABCA2/5* and *ABCB4/6*, in agreement with our previous study where we investigated the expression of transporters in the HCE-T cells [51]. Collectively, our results indicate the expressional activation of genes related to fatty acid oxidation in corneal epithelial cells during cell maturation.

## 5. Conclusion

In conclusion, we propose here a simplified, yet efficient method, SSM-CI for the generation of corneal epithelial cells from hPSCs. SSM-CI provides the advantage of minimizing cell culturing steps using only two defined xenobiotic-free and serum-free culturing mediums in combination with TGF-β and Wnt/β-catenin pathway signaling inhibitors, and human bFGF for 20-25 days (Fig. 5). Compared to both conventional HCE-T cells and hPCEpCs, K1-CEpCs generated by SSM-CI expressed high levels of major differentiation factors and maturation markers. RNA-seq analysis emphasized the successful conversion of hPSCs into the corneal epithelium lineage, including the emergence of corneal progenitor phenotypes prior to cell maturation. Furthermore, we uncovered that despite the requirement for the initial inhibition of TGF-β and Wnt/β-catenin, activation of these pathways is required during the later maturation phase to drive the maturation corneal epithelial cells. Finally, we report an alteration in gene signatures associated with the metabolic characteristics of mature corneal epithelial cells compared to embryonic stem cells, where we observed a decrease in glycolysis-related transcripts and an increase in fatty acid oxidation-related transcripts. This was further attributed to the overexpression of metabolic enzymes and transporter-related transcripts which are mainly responsible for the metabolism of fatty acids.

**Fig. 5.**
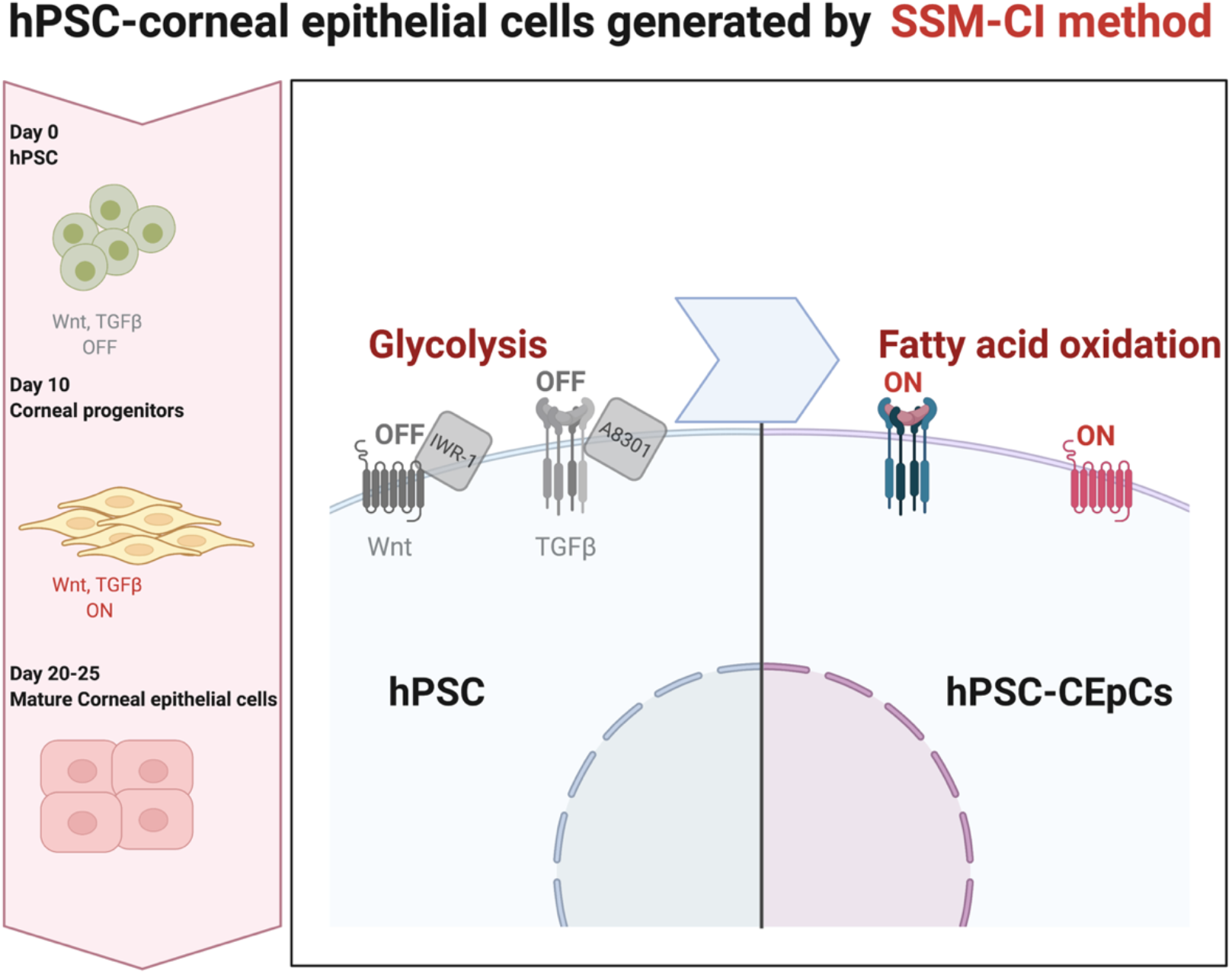
Summary illustration of the SSM-CI method. Xenobiotic-free and serum-free culturing method in combination with chemical inhibitors of the TGF-β pathway, Wnt/β-catenin signaling, and human bFGF for 25 days. The initial inhibition of TGF-β and Wnt/β-catenin at the early stage of differentiation (SWICH OFF) followed by SWITCH ON at the late maturation stage. An alteration in gene signatures associated with the metabolic characteristics of mature corneal epithelial cells: a decrease in glycolysis-related transcripts and an increase in fatty acid oxidation-related transcripts. Created with BioRender.com

## Ethics approval

The human embryonic stem cells were used following the guidelines of the ethics committee of Kyoto University (Approved number: ES3-9).

## Availability of data and material

RNA-seq data have been deposited to the GEO repository under GEO accession number GSE193514.

## Authors Contributions

R. A. conceptualized and managed the project, designed, and performed experiments, analyzed and interpreted data, visualized the data, and wrote the manuscript. K. K. contributed to the experimental design, data interpretation, data visualization, and editing of the manuscript. All authors critically reviewed the manuscript and agreed with the publication.

## Acknowledgments

We acknowledge the WPI-iCeMS supported by the World Premier International Research Centre Initiative (WPI), MEXT, Japan. And the Ritsumeikan Global Innovation Research Organization (R-GIRO).

## References

1. Reichl, S. Cell culture models of the human cornea - a comparative evaluation of their usefulness to determine ocular drug absorption in-vitro. J. Pharm. Pharmacol. 2008, 60, 299–307. doi:10.1211/jpp.60.3.0004.

2. Niederkorn, J.Y.; Meyer, D.R.; Ubelaker, J.E.; Martin, J.H. Ultrastructural and immunohistological characterization of the SIRC corneal cell line. Vitr. Cell. Dev. Biol. - Anim. 1990, 26, 923–930. doi:10.1007/BF02624618.

3. Araki-Sasaki, K.; Ohashi, Y.; Sasabe, T.; Hayashi, K.; Watanabe, H.; Tano, Y.; Handa, H. An SV40-immortalized human corneal epithelial cell line and its characterization. Invest. Ophthalmol. Vis. Sci. 1995, 36, 614–621.

4. Lorsch, J.R.; Collins, F.S.; Lippincott-Schwartz, J. Fixing problems with cell lines. Science. 2014, 346, 1452–1453.

5. Rubelowski, A.K.; Latta, L.; Katiyar, P.; Stachon, T.; Käsmann-Kellner, B.; Seitz, B.; Szentmáry, N. HCE-T cell line lacks cornea-specific differentiation markers compared to primary limbal epithelial cells and differentiated corneal epithelium. Graefe’s Arch. Clin. Exp. Ophthalmol. 2020, 258, 565–575. doi:10.1007/s00417-019-04563-0.

6. Di Girolamo, N.; Chow, S.; Richardson, A.; Wakefield, D. Contamination of primary human corneal epithelial cells with an SV40-transformed human corneal epithelial cell line: A lesson for cell biologists in good laboratory practice. Investig. Ophthalmol. Vis. Sci. 2016, 57, 611–616.

7. Geraghty, R.J.; Capes-Davis, A.; Davis, J.M.; Downward, J.; Freshney, R.I.; Knezevic, I.; Lovell-Badge, R.; Masters, J.R.W.; Meredith, J.; Stacey, G.N.; et al. Guidelines for the use of cell lines in biomedical research. Br. J. Cancer. 2014, 111, 1021–1046.

8. Kitagawa, K.; Kojima, M.; Sasaki, H.; Shui, Y.B.; Chew, S.J.; Cheng, H.M.; Ono, M.; Morikawa, Y.; Sasaki, K. Prevalence of primary cornea guttata and morphology of corneal endothelium in aging Japanese and Singaporean subjects. Ophthalmic Res. 2002, 34, 135–138. doi:10.1159/000063656.

9. Kahn, C.R.; Young, E.; Ihn Hwan Lee Rhim, J.S. Human corneal epithelial primary cultures and cell lines with extended life span: In vitro model for ocular studies. Investig. Ophthalmol. Vis. Sci. 1993, 34, 3429–3441.

10. Hayashi, R.; Ishikawa, Y.; Katori, R.; Sasamoto, Y.; Taniwaki, Y.; Takayanagi, H.; Tsujikawa, M.; Sekiguchi, K.; Quantock, A.J.; Nishida, K. Coordinated generation of multiple ocular-like cell lineages and fabrication of functional corneal epithelial cell sheets from human iPS cells. Nat. Protoc. 2017, 12, 683–696. doi:10.1038/nprot.2017.007.

11. Mikhailova, A.; Ilmarinen, T.; Uusitalo, H.; Skottman, H. Small-Molecule Induction Promotes Corneal Epithelial Cell Differentiation from Human Induced Pluripotent Stem Cells. Stem Cell Reports 2014, 2, 219–231, doi:10.1016/j.stemcr.2013.12.014.

12. Brzeszczynska, J.; Samuel, K.; Greenhough, S.; Ramaesh, K.; Dhillon, B.; Hay, D.C.; Ross, J.A. Differentiation and molecular profiling of human embryonic stem cell-derived corneal epithelial cells. Int. J. Mol. Med. 2014, 33, 1597–1606. doi:10.3892/ijmm.2014.1714.

13. Yang, J.; Park, J.W.; Zheng, D.; Xu, R.H. Universal corneal epithelial-like cells derived from human embryonic stem cells for cellularization of a corneal scaffold. Transl. Vis. Sci. Technol. 2018, 7, 23. doi:10.1167/tvst.7.5.23.

14. Kumagai, H.; Suemori, H.; Uesugi, M.; Nakatsuji, N.; Kawase, E. Identification of small molecules that promote human embryonic stem cell self-renewal. Biochem. Biophys. Res. Commun. 2013, 434, 710–716. doi:10.1016/j.bbrc.2013.03.061.

15. Carpenter, A.E.; Jones, T.R.; Lamprecht, M.R.; Clarke, C.; Kang, I.H.; Friman, O.; Guertin, D.A.; Chang, J.H.; Lindquist, R.A.; Moffat, J.; et al. CellProfiler: image analysis software for identifying and quantifying cell phenotypes. Genome Biol. 2006, 7, R100, doi:10.1186/gb-2006-7-10-r100.

16. Bolger, A.M.; Lohse, M.; Usadel, B. Trimmomatic: A flexible trimmer for Illumina sequence data. Bioinformatics 2014, 30, 2114–2120. doi:10.1093/bioinformatics/btu170.

17. Kim, D.; Langmead, B.; Salzberg, S.L. HISAT: a fast spliced aligner with low memory requirements. Nat. Methods 2015, 12, 357–360. doi:10.1038/nmeth.3317.

18. Li, H.; Handsaker, B.; Wysoker, A.; Fennell, T.; Ruan, J.; Homer, N.; Marth, G.; Abecasis, G.; Durbin, R. The Sequence Alignment/Map format and SAMtools. Bioinformatics 2009, 25, 2078–2090. doi:10.1093/bioinformatics/btp352.

19. Pertea, M.; Pertea, G.M.; Antonescu, C.M.; Chang, T.C.; Mendell, J.T.; Salzberg, S.L. StringTie enables improved reconstruction of a transcriptome from RNA-seq reads. Nat. Biotechnol. 2015, 33, 290–295. doi:10.1038/nbt.3122.

20. Raudvere, U.; Kolberg, L.; Kuzmin, I.; Arak, T.; Adler, P.; Peterson, H.; Vilo, J. G:Profiler: A web server for functional enrichment analysis and conversions of gene lists (2019 update). Nucleic Acids Res. 2019, 47, W191–W198. doi:10.1093/nar/gkz369.

21. Renesh Bedre Bioinformatics data analysis and visualization toolkit. Zenodo 2020, doi:10.5281/zenodo.3965241.

22. Schinzel, R.T.; Ahfeldt, T.; Lau, F.H.; Lee, Y.K.; Cowley, A.; Shen, T.; Peters, D.; Lum, D.H.; Cowan, C.A. Efficient culturing and genetic manipulation of human pluripotent stem cells. PLoS One 2011, 6, e27495. doi:10.1371/journal.pone.0027495.

23. Chen, G.; Gulbranson, D.R.; Hou, Z.; Bolin, J.M.; Ruotti, V.; Probasco, M.D.; Smuga-Otto, K.; Howden, S.E.; Diol, N.R.; Propson, N.E.; et al. Chemically defined conditions for human iPSC derivation and culture. Nat. Methods 2011, 8, 424–429. doi:10.1038/nmeth.1593.

24. Kato, R.; Matsumoto, M.; Sasaki, H.; Joto, R.; Okada, M.; Ikeda, Y.; Kanie, K.; Suga, M.; Kinehara, M.; Yanagihara, K.; et al. Parametric analysis of colony morphology of non-labelled live human pluripotent stem cells for cell quality control. Sci. Rep. 2016, 6, 34009. doi:10.1038/srep34009.

25. Lippmann, E.S.; Estevez-Silva, M.C.; Ashton, R.S. Defined human pluripotent stem cell culture enables highly efficient neuroepithelium derivation without small molecule inhibitors. Stem Cells 2014, 32, 1032–1042. doi:10.1002/stem.1622.

26. Hollmann, E.K.; Bailey, A.K.; Potharazu, A. V.; Neely, M.D.; Bowman, A.B.; Lippmann, E.S. Accelerated differentiation of human induced pluripotent stem cells to blood-brain barrier endothelial cells. Fluids Barriers CNS 2017, 14, doi:10.1186/s12987-017-0059-0.

27. Folmes, C.D.L.; Nelson, T.J.; Martinez-Fernandez, A.; Arrell, D.K.; Lindor, J.Z.; Dzeja, P.P.; Ikeda, Y.; Perez-Terzic, C.; Terzic, A. Somatic oxidative bioenergetics transitions into pluripotency-dependent glycolysis to facilitate nuclear reprogramming. Cell Metab. 2011, 14, 264–271. doi:10.1016/j.cmet.2011.06.011.

28. Takubo, K.; Nagamatsu, G.; Kobayashi, C.I.; Nakamura-Ishizu, A.; Kobayashi, H.; Ikeda, E.; Goda, N.; Rahimi, Y.; Johnson, R.S.; Soga, T.; et al. Regulation of glycolysis by Pdk functions as a metabolic checkpoint for cell cycle quiescence in hematopoietic stem cells. Cell Stem Cell 2013, 12, 49–61. doi:10.1016/j.stem.2012.10.011.

29. Shyh-Chang, N.; Daley, G.Q.; Cantley, L.C. Stem cell metabolism in tissue development and aging. Dev. 2013, 140, 2535–2547.

30. Oosterveen, T.; Garção, P.; Moles-Garcia, E.; Soleilhavoup, C.; Travaglio, M.; Sheraz, S.; Peltrini, R.; Patrick, K.; Labas, V.; Combes-Soia, L.; et al. Pluripotent stem cell derived dopaminergic subpopulations model the selective neuron degeneration in Parkinson’s disease. Stem Cell Reports 2021, 16, 2718 –2735. doi:10.1016/j.stemcr.2021.09.014.

31. Muckom, R.; Bao, X.; Tran, E.; Chen, E.; Murugappan, A.; Dordick, J.S.; Clark, D.S.; Schaffer, D. V. High-throughput 3D screening for differentiation of hPSC-derived cell therapy candidates. Sci. Adv. 2020, 6, eaaz1457. doi:10.1126/sciadv.aaz1457.

32. Lian, X.; Zhang, J.; Azarin, S.M.; Zhu, K.; Hazeltine, L.B.; Bao, X.; Hsiao, C.; Kamp, T.J.; Palecek, S.P. Directed cardiomyocyte differentiation from human pluripotent stem cells by modulating Wnt/β-catenin signaling under fully defined conditions. Nat. Protoc. 2013, 8, 162–175. doi:10.1038/nprot.2012.150.

33. Theerakittayakorn, K.; Nguyen, H.T.; Musika, J.; Kunkanjanawan, H.; Imsoonthornruksa, S.; Somredngan, S.; Ketudat-Cairns, M.; Parnpai, R. Differentiation induction of human stem cells for corneal epithelial regeneration. Int. J. Mol. Sci. 2020, 21, 7834.

34. Chavali, V.R.M.; Haider, N.; Rathi, S.; Vrathasha, V.; Alapati, T.; He, J.; Gill, K.; Nikonov, R.; Duong, T.T.; McDougald, D.S.; et al. Dual SMAD inhibition and Wnt inhibition enable efficient and reproducible differentiations of induced pluripotent stem cells into retinal ganglion cells. Sci. Rep. 2020, 10, 11828. doi:10.1038/s41598-020-68811-8.

35. Takamiya, M.; Stegmaier, J.; Kobitski, A.Y.; Schott, B.; Weger, B.D.; Margariti, D.; Delgado, A.R.C.; Gourain, V.; Scherr, T.; Yang, L.; et al. Pax6 organizes the anterior eye segment by guiding two distinct neural crest waves. PLoS Genet. 2020, 16, e1008774. doi:10.1371/journal.pgen.1008774.

36. Tojo, M.; Hamashima, Y.; Hanyu, A.; Kajimoto, T.; Saitoh, M.; Miyazono, K.; Node, M.; Imamura, T. The ALK-5 inhibitor A-83-01 inhibits Smad signaling and epithelial-to-mesenchymal transition by transforming growth factor-β. Cancer Sci. 2005, 96, 791–800. doi:10.1111/j.1349-7006.2005.00103.x.

37. Callaerts, P.; Halder, G.; Gehring, W.J. Pax-6 in development and evolution. Annu. Rev. Neurosci. 1997, 20, 483–532.

38. Gehring, W.J.; Ikeo, K. Pax 6: Mastering eye morphogenesis and eye evolution. Trends Genet. 1999, 15, 371–377.

39. Zhang, J.; Upadhya, D.; Lu, L.; Reneker, L.W. Fibroblast growth factor receptor 2 (FGFR2) is required for corneal epithelial cell proliferation and differentiation during embryonic development. PLoS One 2015, 10, e0117089. doi:10.1371/journal.pone.0117089.

40. Li, W.; Chen, Y.T.; Hayashida, Y.; Blanco, G.; Kheirkah, A.; He, H.; Chen, S.Y.; Liu, C.Y.; Tseng, S.C.G. Down-regulation of Pax6 is associated with abnormal differentiation of corneal epithelial cells in severe ocular surface diseases. J. Pathol. 2008, 214, 114–122. doi:10.1002/path.2256.

41. Liu, C.Y.; Zhu, G.; Westerhausen-Larson, A.; Converse, R.; Candace, W.C.K.; Sun, T.T.; Winston, W.Y.K. Cornea-specific expression of k12 keratin during mouse development. Curr. Eye Res. 1993, 12, 963–974. doi:10.3109/02713689309029222.

42. Merjava, S.; Brejchova, K.; Vernon, A.; Daniels, J.T.; Jirsova, K. Cytokeratin 8 is expressed in human corneoconjunctival epithelium, particularly in limbal epithelial cells. Investig. Ophthalmol. Vis. Sci. 2011, 52, 787–794. doi:10.1167/iovs.10-5489.

43. Kao, W.W.Y. Keratin expression by corneal and limbal stem cells during development. Exp. Eye Res. 2020, 200, 108206.

44. Waters, J.M.; Richardson, G.D.; Jahoda, C.A.B. Keratin 10 (K10) is expressed suprabasally throughout the limbus of embryonic and neonatal rat corneas, with interrupted expression in the adult limbus. Exp. Eye Res. 2009, 89, 435–438. doi:10.1016/j.exer.2009.03.018.

45. Li, D.Q.; Lokeshwar, B.L.; Solomon, A.; Monroy, D.; Ji, Z.; Pflugfelder, S.C. Regulation of MMP-9 production by human corneal epithelial cells. Exp. Eye Res. 2001, 73, 449–459. doi:10.1006/exer.2001.1054.

46. Blalock, T.D.; Spurr-Michaud, S.J.; Tisdale, A.S.; Heimer, S.R.; Gilmore, M.S.; Ramesh, V.; Gipson, I.K. Functions of MUC16 in corneal epithelial cells. Investig. Ophthalmol. Vis. Sci. 2007, 48, 4509–4518. doi:10.1167/iovs.07-0430.

47. Arkell, R.M.; Tam, P.P.L. Initiating head development in mouse embryos: Integrating signalling and transcriptional activity. Open Biol. 2012, 2, 120030. doi:10.1098/rsob.120030.

48. Warburg, O.; Wind, F.; Negelein, E. The metabolism of tumors in the body. J. Gen. Physiol. 1927, 8, 519–530. doi:10.1085/jgp.8.6.519.

49. Ito, K.; Suda, T. Metabolic requirements for the maintenance of self-renewing stem cells. Nat. Rev. Mol. Cell Biol. 2014, 15, 243–256.

50. Anderson, C.M.; Stahl, A. SLC27 fatty acid transport proteins. Mol. Aspects Med. 2013, 34, 516–528.

51. Abdalkader, R.; Chaleckis, R.; Wheelock, C.E.; Kamei, K. ichiro Spatiotemporal determination of metabolite activities in the corneal epithelium on a chip. Exp. Eye Res. 2021, 209, 108646. doi:10.1016/j.exer.2021.108646.

