## Supplementary Information for "An efficient simplified method for the generation of corneal epithelial cells from human pluripotent stem cells"

*Corresponding authors:

**Supplementary Table 1. List of differentiation reagents and mediums**

| Compound | Solvent | Maker | Catalog number | Final conc. |
|---|---|---|---|---|
| IWR-1 endo | DMSO | Selleck | S7086 | 0.3-10 μM |
| A83-01 | DMSO | Wako Fujifilm | 039-24111 | 0.3-10 μM |
| bFGF | D.H2O | Wako Fujifilm | 064-04541 | 50 ng $mL^{-1}$ |
| Essential 8 Medium | NA | Thermo Fisher | A1517001 | NA |
| Essential 6 Medium | NA | Thermo Fisher | A1516401 | NA |

NA: not applicable

### Supplementary Table 2. List of antibodies

| Target protein | Host species | Maker | Catalog number | Ratio (v/v) |
|---|---|---|---|---|
| CK12 | Mouse | Santacruz | sc-515882 | 1:200 |
| PAX6 | Rabbit | Abcam | AB5790-100 | 1:200 |

**Supplementary Figure 1**

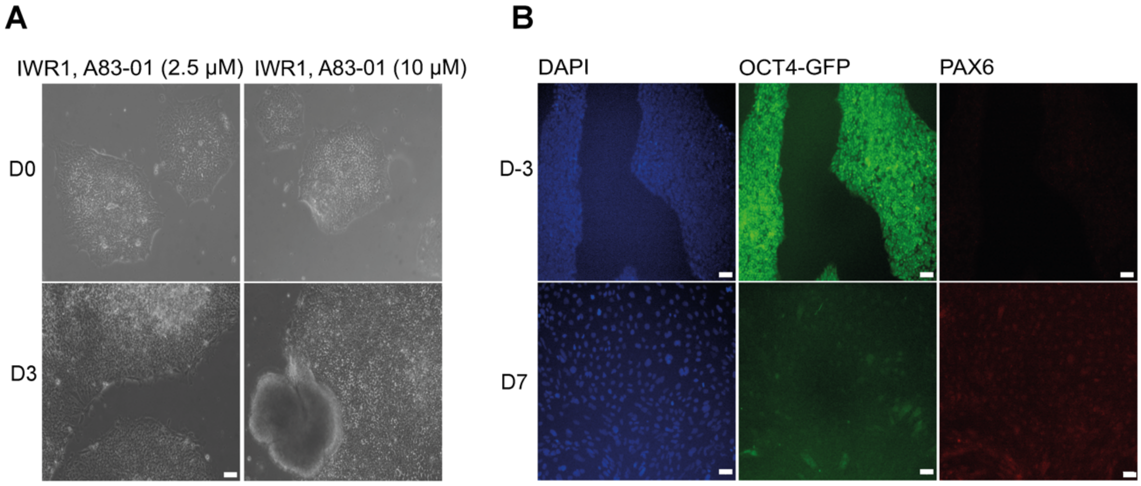


**Fig. S1. Optimization of the concentration of chemical inhibitors**.

Human ES cells with enhanced green fluorescent protein (EGFP) driven by *the OCT4* promoter (K1-OCT4-EGFP) were treated with IWR-1 endo and A83-01 at concentrations of 2.5% and 10 μM, respectively (A). Bright-field images of cells on days 0 (D0) and 3(D3). (B). Fluorescent micrograph images indicating the expression of OCT4-EGFP and the eye developmental marker PAX6. Blue: DAPI, green: OCT4, red: PAX6 scale bar, 50 μm.

**Supplementary Figure 2**

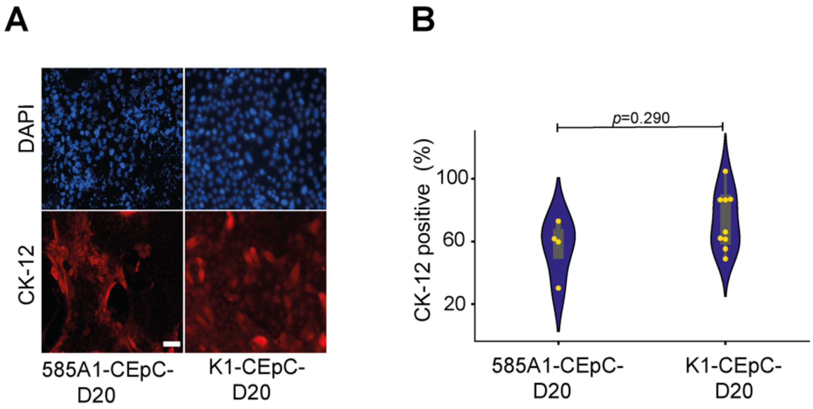


**Fig. S2. Positive expression of CK-12 in ES (K1)/ iPS(585A1) derived corneal epithelial cells**.

(A). Fluorescent micrograph images indicating the expression of the corneal maturation marker (CK1-2). Blue: DAPI, Red: CK12. Scale bar, 50 µm. (B). Violin plot of indicating the percentage of CK-12 positive cells. Blue: DAPI, Green: OCT4, Red: PAX6 Scale bar, 50 µm. The *p*-values were determined using the unpaired t-test.

### Supplementary Figure 3

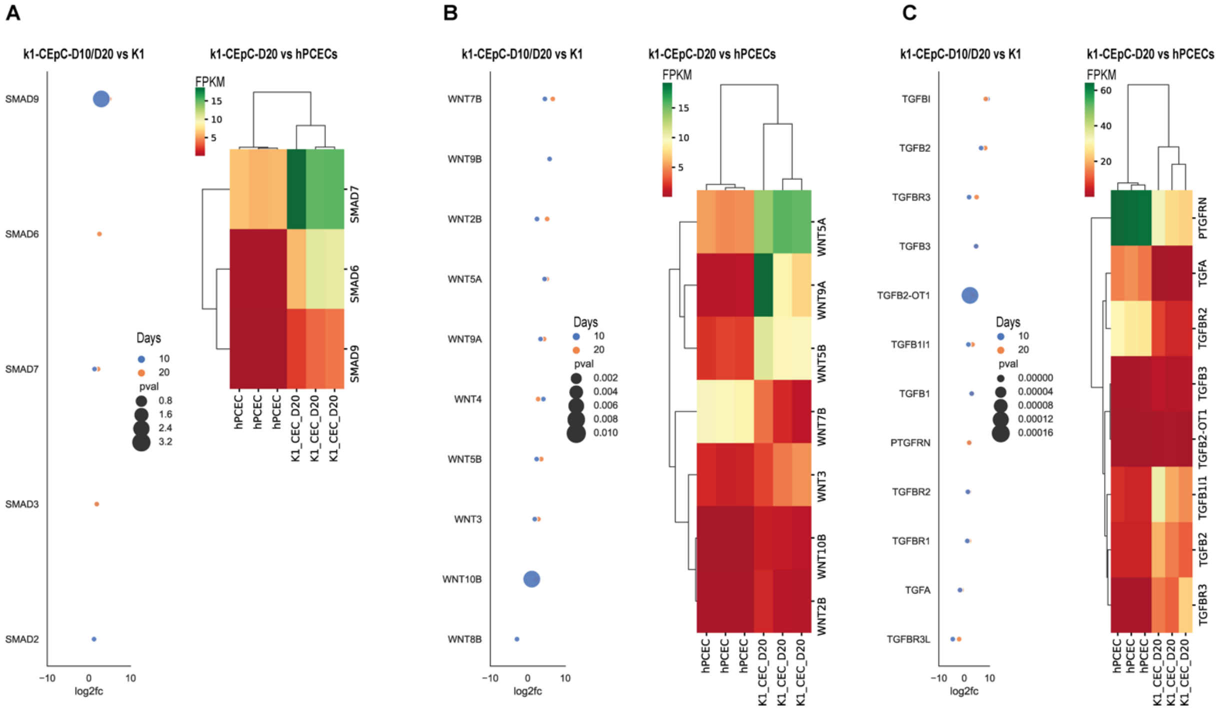


**Fig. S3. The expression of TGF-β and Wnt/β-catenin related genes**

(A). SMADs family (B). WNT family. (C). TGF-β family. Right: DEGs at D10 (blue) and D20 (orange). Genes with 2-fold change and a p-value <0.05 were applied. Left: Heatmap of selected genes in K1, hPCEpC, and K1-CEpCs-D20.

### Supplementary Figure 4

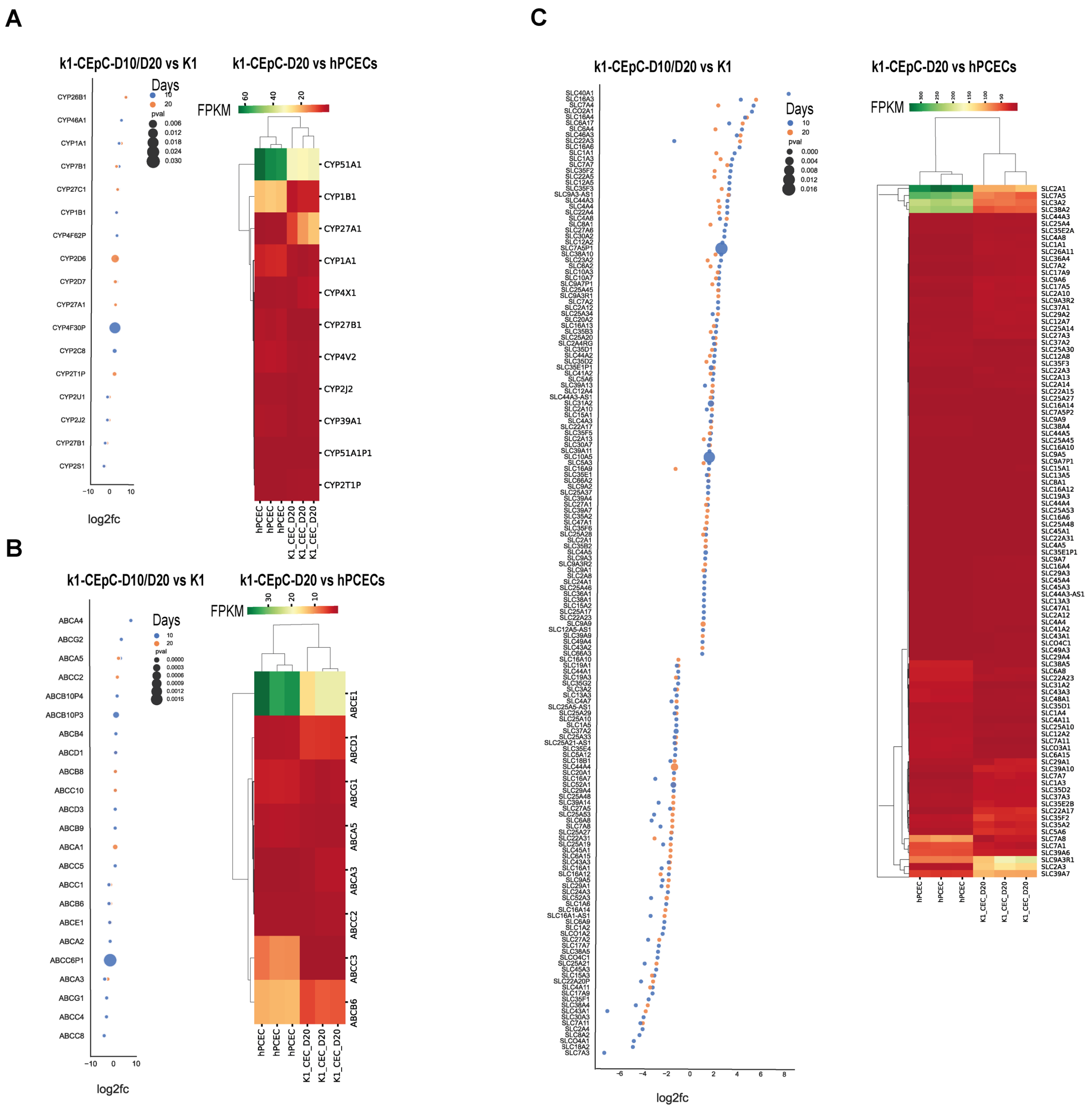


**Fig. S4. The expression of metabolic enzymes and transporters-related genes**

(A). CYPs family (B). ABC transporters family. (C). SLC transporters family. Right: DEGs at D10 (blue) and D20 (orange). Genes with 2-fold change and a p-value <0.05 were applied. Left: Heatmap of selected genes in K1, hPCEpC, and K1-CEpCs-D20.
